# Short-chain fatty acid production by gut microbiota from children with obesity is linked to bacterial community composition and prebiotic choice

**DOI:** 10.1101/2020.04.10.035808

**Authors:** Zachary C. Holmes, Justin D. Silverman, Holly K. Dressman, Zhengzheng Wei, Eric P. Dallow, Sarah C. Armstrong, Patrick C. Seed, John F. Rawls, Lawrence A. David

## Abstract

Pediatric obesity remains a public health burden and continues to increase in prevalence. The gut microbiota plays a causal role in obesity and is a promising therapeutic target. Specifically, the microbial production of short-chain fatty acids (SCFA) from the fermentation of otherwise indigestible dietary carbohydrates may protect against pediatric obesity and metabolic syndrome. Still, it has not been demonstrated that therapies involving microbiota-targeting carbohydrates, known as prebiotics, will enhance gut bacterial SCFA production in children and adolescents with obesity (age 10-18). Here, we used an *in vitro* system to examine the SCFA production by fecal microbiota from 17 children with obesity when exposed to five different commercially available over-the-counter (OTC) prebiotic supplements. We found microbiota from all 17 patients actively metabolized most prebiotics. Still, supplements varied in their acidogenic potential. Significant inter-donor variation also existed in SCFA production, which 16S rRNA sequencing supported as being associated with differences in the host microbiota composition. Last, we found that neither fecal SCFA concentration, microbiota SCFA production capacity, nor markers of obesity positively correlated with one another. Together, these *in vitro* findings suggest the hypothesis that OTC prebiotic supplements may be unequal in their ability to stimulate SCFA production in children and adolescents with obesity, and that the most acidogenic prebiotic may differ across individuals.

**IMPORTANCE:** Pediatric obesity remains a major public health problem in the US, where 17% of children and adolescents are obese, and rates of pediatric ‘severe obesity’ are increasing. Children and adolescents with obesity face higher health risks, and non-invasive therapies for pediatric obesity often have limited success. The human gut microbiome has been implicated in adult obesity, and microbiota-directed therapies can aid weight loss in adults with obesity. However, less is known about the microbiome in *pediatric* obesity, and microbiota-directed therapies are understudied in children and adolescents. Our research has two important findings: 1) dietary prebiotics (fiber) cause the microbiota from adolescents with obesity to produce more SCFA, and 2) the effectiveness of each prebiotic is donor-dependent. Together, these findings suggest that prebiotic supplements could help children and adolescents with obesity, but that these therapies may not be one-size-fits-all.

## Introduction

Approximately 17% of children in the United States have obesity, and the prevalence continues to increase among all ages and populations (1). The prevalence of pediatric obesity is even higher in Hispanic and African American populations in the United States, where rates of severe obesity continue to increase (1). Children with obesity have an increased risk of adverse health events and incur higher healthcare costs (2–4). Despite the severity of the pediatric obesity epidemic, current common treatment strategies centered around lifestyle changes, including behavioral, dietary, and exercise interventions, often fail or have limited success (5). The high prevalence of pediatric obesity, coupled with the low success rate of common interventions, highlights the need for more efficacious, safe strategies to lower BMI in children and adolescents.

The human gut microbiome has emerged as a promising therapeutic target in pediatric obesity. Over the past decade, differences in gut microbial community composition and metabolic activity between obese and lean individuals have been observed (6–8). Causal links have also been established; fecal transplantation can transfer the obesity phenotype from obese donors to lean recipients and recapitulate some key metabolic changes in human obesity (9). Multiple mechanisms for this link have been proposed, including increased energy harvest by obese microbiota (10), activation of enteroendocrine signaling pathways by SCFA (11–13), modulation of glucose and energy homeostasis through bile acid signaling (14), and increased local and systemic inflammation caused by a variety of microbial metabolites (15).

Recent attention in obesity research has been specifically drawn to the role of microbially-derived short-chain fatty acids (SCFA). SCFAs, primarily acetate, propionate, and butyrate, are produced by enteric microbes as end products of anaerobic fermentation of undigested, microbially-accessible dietary carbohydrates, and serve a variety of important roles in the gut. Of particular interest is the SCFA butyrate, which serves as the primary nutrient source for colonocytes (16) and functions as a histone deacetylase inhibitor (17, 18). Through its inhibition of NF-κB signaling in colonocytes, butyrate contributes to barrier integrity maintenance and reduces levels of intestinal inflammation markers (19–22). Acetate, propionate, and butyrate also each activate G-protein coupled receptors (GPR) that modulate key metabolic hormones including peptide YY (PYY) and GLP-1 (12, 23). Consistent with these mechanistic findings, mouse studies have shown that supplementation with acetate, propionate, butyrate, or some mixture of these can protect against weight gain, improve insulin sensitivity, and reduce obesity-associated inflammation (24–29). Given the experimental evidence for SCFA supplementation having an anti-obesogenic effect in a murine system, maintaining high levels of SCFA during a weight loss treatment may improve results (27).

If increasing SCFA levels is a potential approach to promote weight loss in children, prebiotic supplementation may provide an effective and low-risk adjunctive therapy. Prebiotics are dietary carbohydrates that are indigestible by human-produced enzymes and thus survive transit to the lower GI tract. Once in the colon, prebiotics serve as carbon sources for bacterial fermentation, which in turn yield SCFAs as metabolic end products (30, 31). Multiple types of prebiotics (*e.g*. fructooligosaccharides (FOS), and inulin-type fructans) have been tested in children with obesity ranging from ages 7-18. In select cases, these treatments have been associated with smaller increases in BMI and fat mass (32), and reductions in body weight z-scores, body fat, and trunk fat (33). Still, other prebiotic trials in overweight children have reported no significant beneficial effects (34).

Interpreting the mixed outcomes of prior prebiotic clinical trials in pediatric obesity though is complicated by several challenges. First, *in vivo* studies in pediatric obesity to date have each used only one prebiotic supplement due to the logistical constraints of clinical trials (32–34). Trials employing testing only a single type of supplement hinder the ability to generalize conclusions regarding the efficacy of prebiotics and also make it challenging to determine whether some prebiotics are inherently more acidogenic than others. Second, *in vivo* trials in healthy adults have shown substantial inter-individual variation in the single prebiotic effects on stool SCFA concentration (30, 31, 35). Variation in the primary and secondary outcomes could be due to differences in microbial SCFA production; or differences in host physiology, such as SCFA absorption potential. Third, while SCFA concentrations have been shown to be altered in children with overweight or obesity (36), changes in fecal SCFA during dietary intervention have not been measured in past *in vivo* studies in pediatric populations. If prebiotics mediate their effects through SCFA (33, 34, 37), directly tracking SCFAs could help determine treatment success. Fourth, *in vivo* studies in adults, especially those with obesity, may be confounded by the concurrence of chronic disease and the medications a person may be taking to treat chronic disease.

In this study, we have taken an *in vitro* approach to address the limitations of prior human studies. An *in vitro* approach facilitates more direct comparisons of different prebiotic supplements: the higher-throughput of *in vitro* experiments allows wider variety of prebiotics to be tested; and, the effects of these supplements can be tested on identical microbiota samples, rather than over time within subjects, which is confounded by microbiota drift over time (38), as well as inconsistencies in dietary composition. Taking an *in vitro* approach to studying the effects of prebiotics on gut microbiota allows a more direct investigation of microbial SCFA production, as we can study the effects of prebiotic supplementation independent of the effects of host absorption (39, 40). Using a preclinical *in vitro* fermentation model, and samples from adolescents with obesity who have not developed long-term complications, we pursued three specific lines of inquiry: 1) whether different types of prebiotics lead to differences in SCFA production by gut microbiota from adolescents with obesity; 2) whether the effects of prebiotics are shaped by inter-individual differences in gut microbiota structure; and, 3) whether fecal SCFA production is associated with protection from obesity.

## Results

### SCFA production capacity

To measure SCFA production by gut microbiota, we adapted the *in vitro* approach of Edwards et al. 1996 (41). This method was specifically designed to study fermentation of starch in the human lower GI tract, and has since been used to measure metabolite production from human stool samples when exposed to prebiotic fiber (42–44). In brief, we homogenized previously frozen feces in reduced phosphate buffered saline (pH 7.0 ± 0.1) to create a fecal slurry with a final concentration of 100g/L (Figure 1). These fecal slurries were then supplied with each of five prebiotic carbon sources, and a carbon-free control, and allowed to ferment at 37°C in anaerobic conditions for 24 hours, to approximate colonic transit time (45). Following the incubation period, the concentrations of SCFA in the samples were measured by gas chromatography. To control for differences in overall cell viability or stool slurry nutrient content between donors, we corrected measurements of SCFA concentration by dividing the treatment SCFA concentration by the control SCFA concentration.

**Figure 1:**
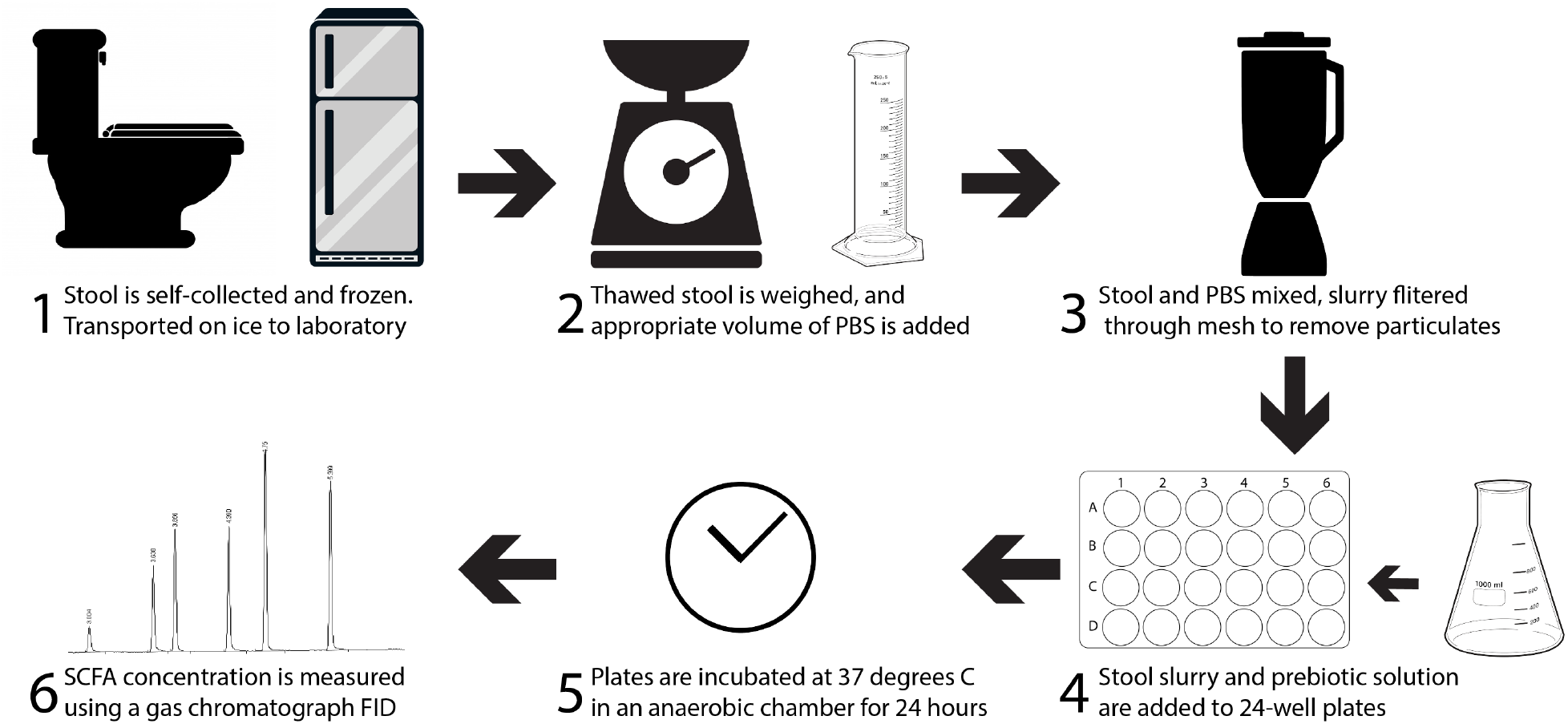
Overview of *in vitro* fermentation methods.

To validate our assay, we ran a series of experiments using feces from validation sample sets. We verified that our control-corrected SCFA production data was not influenced by bacterial abundance (p = 0.38, ρ = 0.14, Spearman correlation; Figure S1). Absolute (not relativized to control) SCFA concentrations are supplied in the supplement (Figures S2 and S3). As our fermentation experiments used previously frozen fecal samples, we verified that total SCFA production was strongly correlated between fresh samples and twice freeze-thawed samples (p < 0.0001, ρ = 0.75, Spearman correlation; Figure S4A). Since we elected to not provide our fermentation reactions with nutrients in excess of what was contained in the fecal slurries, we verified that there existed strong correlation in total SCFA production between PBS-grown and colonic medium-grown cultures (46), both when supplied with dextrin and inulin (Dextrin: p = 0.001, ρ = 0.68; inulin: p = 0.02, ρ = 0.51; Spearman correlations; Figure S5). We found that total SCFA production over control was positively correlated with the pH of starting fecal slurries (p = 0.003, ρ = 0.46; Spearman correlation; Figure S6A). A weaker correlation may exist between SCFA production and the final pH of the fermentation vessels (p = 0.067, ρ = 0.29, Spearman correlation; Figure S6B).

We subsequently applied our assay to fecal microbiota from a cohort of 17 children ranging in age from 10 – 18, Tanner stages 2 – 5, and body-mass index (BMI) from 25.9 – 75.3 (Table 1). We found all 17 individuals demonstrated a net gain of SCFA relative to the control in at least one prebiotic treatment, which led us to conclude that all tested cultures were viable and metabolically active (Figure 2).

**Table 1:** Demographic characteristics of participants in this study. One patient provided samples used in all analyses but was lost to follow-up before providing clinical metadata; that patient is only counted in the total column.

| Variable | Male (n) | Female (n) | BMI Range | Average BMI | Age Range | Average Age | Total |
| --- | --- | --- | --- | --- | --- | --- | --- |
| Value | 6 | 10 | 25.9 – 75.3 | 34.9 | 10-18 | 15.7 | 17 |

**Figure 2:**
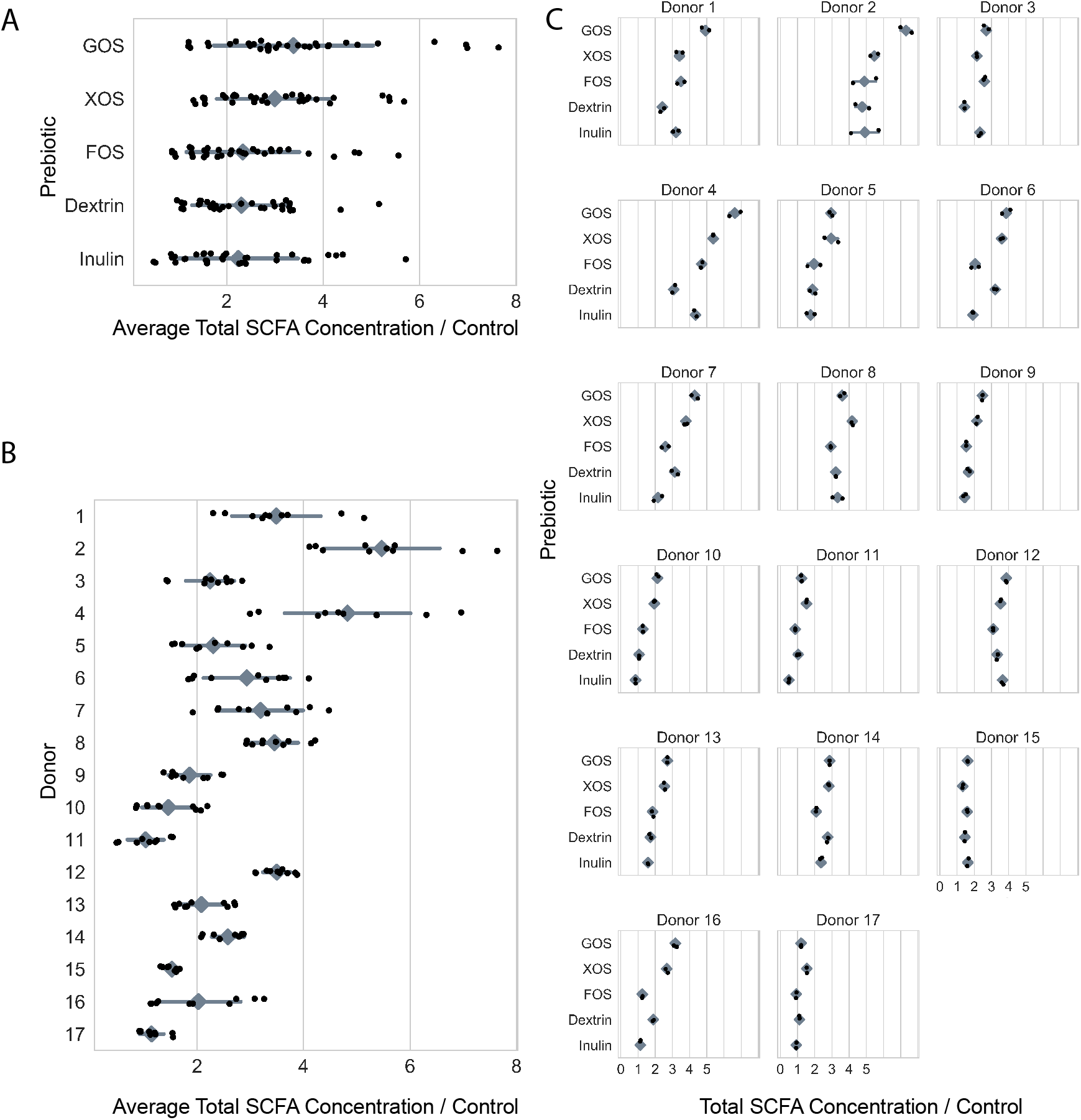
In vitro SCFA production by prebiotic (a), donor (b), and individually (c). In a two-way ANOVA of the effects of ‘Donor’ and ‘Prebiotic’ on ‘SCFA Concentration / Control’, ‘Donor’, ‘Prebiotic’, and their interaction were all statistically significant (p<0.0001, p<0.0001, p<0.0001, respectively). Shown is the total SCFA concentration of an in vitro culture after 24hrs of anaerobic incubation, divided by the SCFA concentration of the corresponding prebiotic-free control culture, for each of five prebiotic growth conditions across 17 donors (black dots). Grey diamonds are means and grey bars are standard deviations. (Absolute SCFA concentrations are depicted in Figure S3).

### Donor and prebiotic both impact SCFA production *in vitro*

We next tested the hypothesis that different prebiotics equally promote the production of SCFA by performing statistical analysis of SCFA production as a function of the prebiotic type and individual identity. Our analysis revealed heterogeneity in the efficacy of prebiotic supplements (two-way ANOVA, p < 0.001; Table S1; Figure 2a), ranging from inulin, which resulted in a 2.35 mean fold change in total SCFA, to GOS, which resulted in 3.55 mean fold change in total SCFA. Frequently, only two or three of the five tested prebiotics resulted in increased total SCFA production within an individual. Our statistical testing also revealed consistent patterns between individuals’ gut microbiota in terms of SCFA production (two-way ANOVA, p < 0.001; Table S1; Figure 2b), with mean fold change in SCFA over control ranging from 2.37 to 6.12. Within individuals, the average fold change in SCFA concentration in the prebiotic treatments often appeared to be driven by a few strongly acidogenic prebiotics. Last, our analysis indicated a significant interaction between prebiotic type and individual identity (two-way ANOVA, p< 0.001; Table S1; Figure 2c). Because our statistical analysis considered technical replicates as separate experimental conditions, this result suggests the presence of consistent prebiotic/individual responses across *in vitro* assay replicate runs – not whether such interactions are consistent within an individual over time.

### SCFA production *in vitro* predicts the abundance of bacteria in the starting culture

If inter-individual differences in gut microbiota mediated responses to prebiotic treatment, we would expect that specific bacterial taxa, which varied between individuals, could also be associated with SCFA production. To evaluate this hypothesis, we used the R package *stray* (47) to create a Bayesian multinomial logistic normal linear regression (*pibble*) model that tested for correlations between *in vitro* SCFA production in response to each prebiotic and 16S rRNA community composition of patient stool used in the fermentations, at the genus level. This analysis revealed that SCFA production from prebiotics was correlated with the relative abundances of 18 different bacterial genera (95% credible interval not covering 0, Figure 3). Of the 13 genera positively associated with SCFA production, 9 are known or likely fiber degraders (48–52), *Akkermansia*, is often observed to increase in abundance after prebiotic treatment (53), and one, *Methanobrevibacter*, an archaeon hydrogenotrophic methanogen, is known to increase the efficiency of carbohydrate metabolism by the microbiota (54) (Table 3). Most genera identified by *stray* were associated with SCFA production in a limited set of prebiotic treatments. One genus, *Lactobacillus*, is positively associated with SCFA production on XOS, but were negatively associated with SCFA production on GOS. Overall, the presence of specific associations between bacterial taxa and different prebiotics supports a model where different individuals vary in their levels of prebiotic degrading gut bacteria.

**Figure 3:**
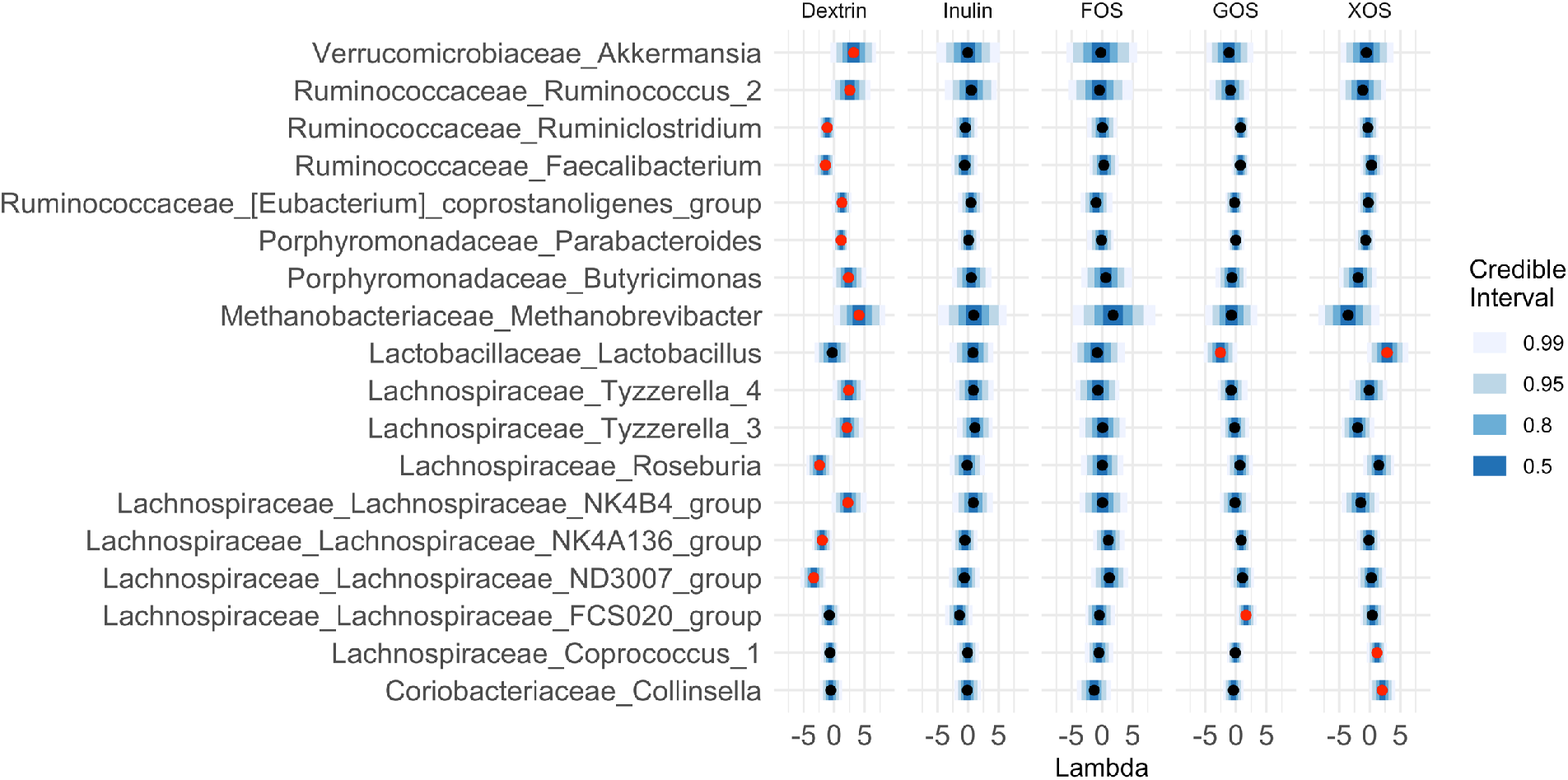
18 genera were found to be credibly associated with SCFA production in at least one of our five prebiotic growth conditions. Shown are the mean Lambda and 99%, 95%, 80%, and 50% credible intervals for all 18 genera credibly associated with at least one prebiotic growth condition, plotted on centered log-ratio (CLR) coordinates. Red centers denote associations with 95% credible intervals that do not cover 0. Lambda represents the strength of the effect of each covariate on each taxa. A lambda of one reflects a unit fold-change in SCFA concentration over control as being associated with a unit fold-change in the CLR-transformed relative abundance of the genus.

### Metrics of obesity do not appear to correlate with SCFA production capacity of stool

Finally, we tested the hypothesis that *in vitro* SCFA production would be associated with obesity-related phenotypes. We compared clinical metadata from individuals, which included BMI, insulin, and HbA1c, with average total SCFA production across prebiotics and found no significant correlations in our population (Spearman correlation; Table 2). Fecal microbial SCFA production capacity may not be directly associated with obesity though because rates of host SCFA uptake likely vary, and this variance may influence host intestinal physiology (55–57). Indeed, in support of the idea that SCFA absorption rate (which was not measured in this study) shape metabolic homeostasis and host health, we observed a negative association between fecal SCFA concentrations and *in vitro* SCFA production across the range of tested prebiotics (Figure 4). Furthermore, if SCFA absorption efficiencies varied by individual, residual fecal SCFA concentrations may not directly reflect the complete effect of bacterial metabolism on obesity. Consistent with this notion, no significant relationships were apparent between concentrations of SCFA in patient stool and clinical markers of obesity measured at enrollment, including BMI, insulin levels, and HbA1c (Table 2), although this may also be explained by uncontrolled patient parameters.

**Figure 4:**
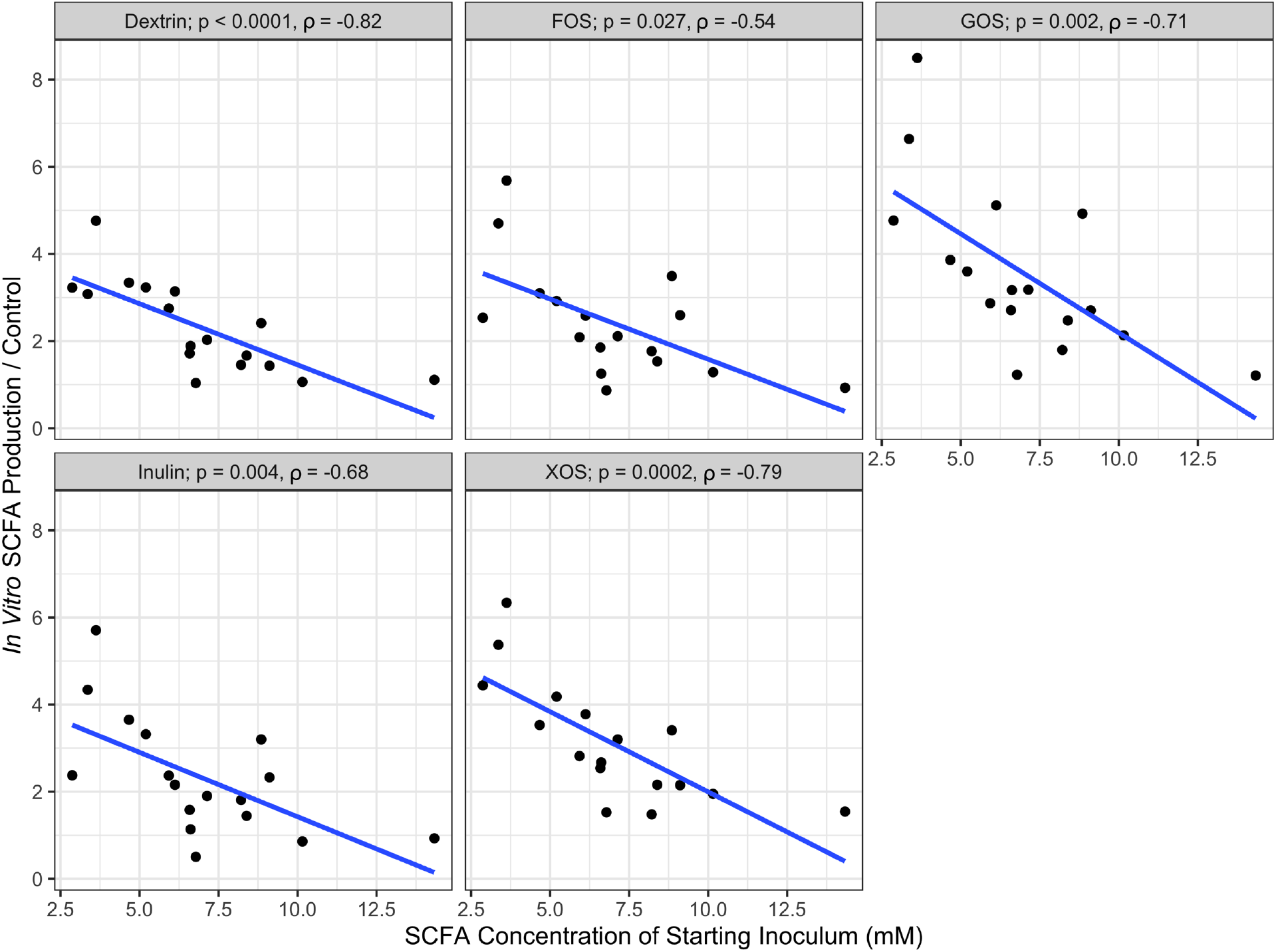
Spearman correlations between *in vitro* SCFA production and SCFA concentration of the starting fecal inoculum. SCFA production is the average of technical replicates, with the linear regression line plotted.

**Table 2:** Neither average SCFA production *in vitro* nor fecal SCFA concentration correlated with metrics of obesity measured in individuals at time of enrollment. P values and ρ from Spearman correlations.

|  | BMI | Insulin | HbA1c |
| --- | --- | --- | --- |
| Average Net SCFA Production | p = 0.98<br>$\rho = -0.007$ | p = 0.63<br>$\rho = 0.13$ | p = 0.75<br>$\rho = 0.083$ |
| Fecal SCFA | p = 0.65<br>$\rho = -0.12$ | p = 0.61<br>$\rho = 0.13$ | p = 0.72<br>$\rho = -0.09$ |

**Table 3:**
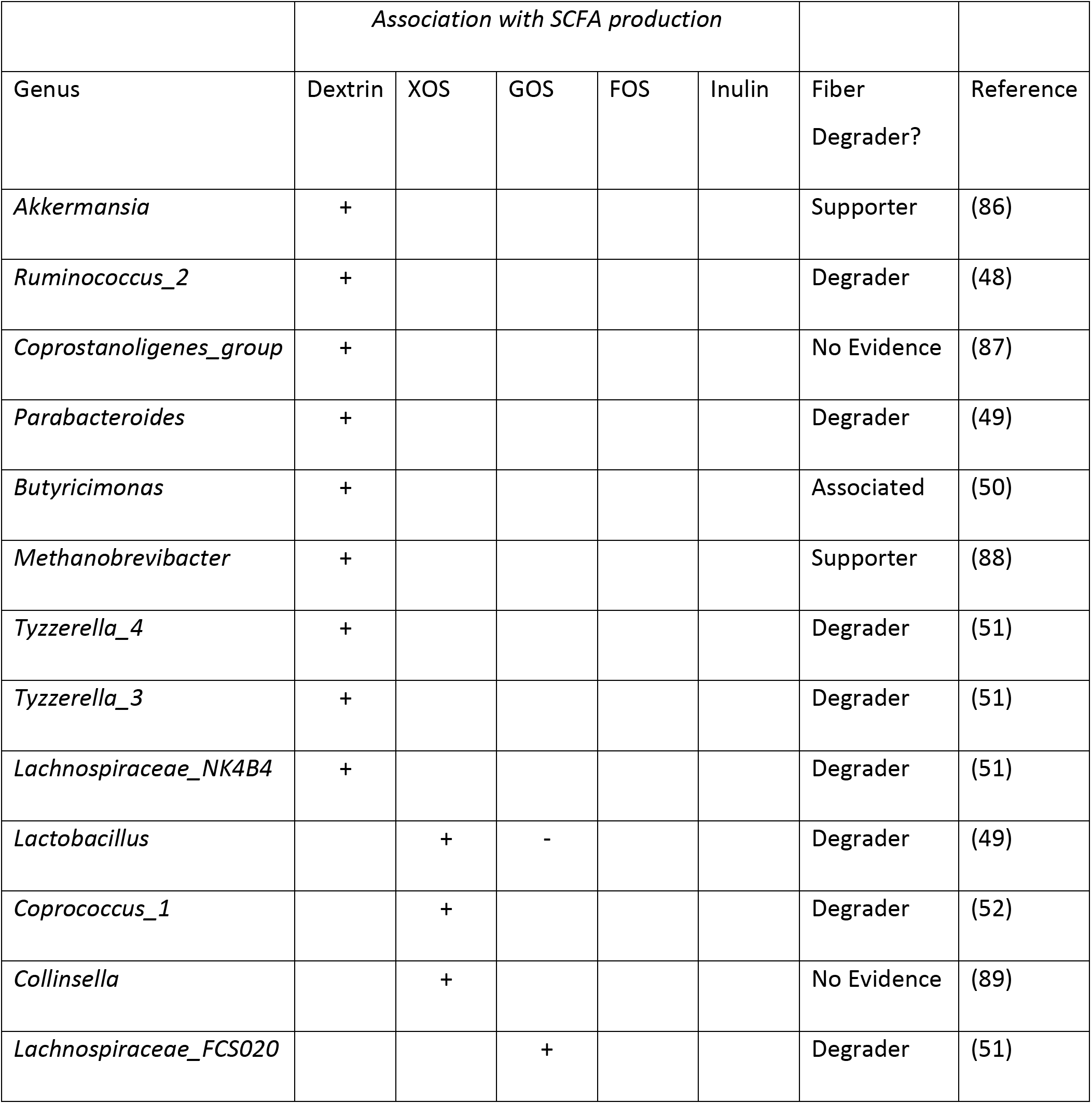
Associations between microbial genera and SCFA production on five different prebiotic substrates.

| Genus | Association with SCFA production |  |  |  |  | Fiber Degrader? | Reference |
| --- | --- | --- | --- | --- | --- | --- | --- |
|  | Dextrin | XOS | GOS | FOS | Inulin |  |  |
| *Akkermansia* | + |  |  |  |  | Supporter | (86) |
| *Ruminococcus_2* | + |  |  |  |  | Degrader | (48) |
| *Coprostanoligenes_group* | + |  |  |  |  | No Evidence | (87) |
| *Parabacteroides* | + |  |  |  |  | Degrader | (49) |
| *Butyricimonas* | + |  |  |  |  | Associated | (50) |
| *Methanobrevibacter* | + |  |  |  |  | Supporter | (88) |
| *Tyzzarella_4* | + |  |  |  |  | Degrader | (51) |
| *Tyzzarella_3* | + |  |  |  |  | Degrader | (51) |
| *Lachnospiraceae_NK4B4* | + |  |  |  |  | Degrader | (51) |
| *Lactobacillus* |  | + | - |  |  | Degrader | (49) |
| *Coprococcus_1* |  | + |  |  |  | Degrader | (52) |
| *Collinsella* |  | + |  |  |  | No Evidence | (89) |
| *Lachnospiraceae_FCS020* |  |  | + |  |  | Degrader | (51) |

## Discussion

In this study we found that the microbiota of all tested adolescents with obesity increased total SCFA production when exposed *in vitro* to at least one prebiotic. Both donor and prebiotic were significant factors in determining SCFA production *in vitro*, as was their interaction. Our modeling revealed distinct associations between specific microbial taxa and SCFA production on different prebiotics. We interpret this result as suggesting that the associated bacteria play a role in the fiber fermenting capacity of the community. We observed no correlations between either stool SCFA concentrations or *in vitro* acidogenic capacity of communities and any metrics of obesity (Table 2).

We have recapitulated previous findings that both donor and prebiotic are important in determining the SCFA production from *in vitro* prebiotic supplementation (31, 50, 58), and we found that not all prebiotics appear equally acidogenic (50). Since our *in vitro* system removes the host as a potential source of variation, our data support a gut microbial role for inter-donor variation in fecal SCFA production. In addition, the strength of the interaction between donor and prebiotic strongly suggests that prebiotics are not one-size-fits-all; rather, inconsistent results from prior studies of prebiotics in pediatric obesity (32, 34, 59) may be due to variation in the SCFA production capacity of individuals’ gut microbiota across the tested prebiotics. Future therapeutic efforts involving prebiotics in patients with obesity may benefit from stratified or personalized treatments. Nutritional therapies that are personalized to individuals’ microbiota are already in development (60).

Murine and *in vitro* studies show that increased signaling through GPRs, mediated by acetate, propionate, and butyrate, increases satiety and insulin sensitivity, while decreasing adipogenesis (12, 23, 61), yet, we did not observe associations between fecal SCFA levels and metrics of obesity. The effects of SCFA on obesity may be masked by uncontrolled patient factors, such as differences in caloric intake and variation in individual nutrient harvest and utilization. In order to observe the effects of SCFA on obesity, it would be necessary to control for these variable physiological and lifestyle parameters, which we did not attempt. These patient factors may also have influenced our inability to observe an association between acidogenic capacity of microbiota and fecal SCFA concentrations. However, this may also be explained by the potential uncoupling of fecal SCFA production and fecal SCFA concentration. *In vitro,* increased luminal concentrations of butyrate have been shown to upregulate the sodium-coupled monocarboxylae transporter SLC5A8 (55), and addition of physiological mixtures of SCFA has been shown to upregulate the monocarboxylate transporter SLC16A1 (62), both of which uptake acetate, propionate, and butyrate from the lumen. Since gut epithelia have the capacity to absorb up to 95% of SCFA before excretion (63), increased host SCFA uptake (triggered by increased gut bacterial production) could, therefore, lead to constant or even decreased fecal SCFA concentrations. This complex relationship could explain the absence of positive correlations we observed between stool SCFA levels and the acidogenic capacity of gut microbiota. It may be necessary to delve further upstream of fecal SCFA concentration by measuring proxies for host SCFA uptakes, such as the expression of SCFA transporters (SLC5A8 and SLC16A1) and SCFA receptors (GPR43, GPR41, and GPR109A) (55).

The primary limitations of this study involve constraints common to *in vitro* culture studies. First, many factors affecting bacterial SCFA production *in vivo* are difficult to replicate *in vitro*, including the availability of nutrients such as nitrogen, the starting concentration of SCFA, the redox state of the environment, and the efficiency of cross-feeding interactions (64, 65). Different metabolic results between prebiotics may have occurred if we provided alternative co-metabolites or nutrients, in addition to the tested prebiotics. We chose our culture conditions, namely a media-free approach that does not add any nutrients beyond what is present in the stool, in an effort to avoid inducing artificial selective conditions within our cultures. Prior experimental digestion studies have shown that prebiotic response patterns can be recapitulated across varying culture conditions (42, 44). Indeed, we found strong correlation in SCFA production between cultures grown with our media-free approach and those grown in a more conventional medium containing added nitrogen, vitamins, minerals, and acetate. Further, this approach allowed us to minimize the influence of the host on measurements of microbiota production of SCFA. We did observe shifts in community composition during the 24 hour fermentations (Figure S7); however, we remained able to find statistical associations between SCFA production capacity and pre-fermentation community composition. A second set of limitations in this study involves our reliance on patient collection of stool. Inter-donor variation in prebiotic response could have originated in technical variation between how patients exposed stool to aerobic conditions (66) or how they froze their samples (67), which in turn could have affected the fraction of viable microbial cells in stool samples. Still, we found a significant correlation between *in vitro* total SCFA production from fresh stool and stool that had been frozen and thawed twice. Variation in donor prebiotic response could also have biological origins due to physiological differences between people (*e.g*. efficiency of food digestion, consistency of stool (68)) or differences in diet, which can lead to variation in stool microbial load and nutrient content (69). Rather than control for a myriad of different sources of variation whose origins we did not measure, we chose the straightforward approach of standardizing donor samples by employing a consistent concentration of stool slurry (5% w/v stool in PBS) in our experiments.

Future work to address these limitations could test multiple stool samples per subject to confirm whether the observed variation in prebiotic response is durable between individuals over time. Future studies could also examine the correlation between the metabolic effects of prebiotic supplementation *in vitro* and *in vivo* using randomized human trials that couple human prebiotic supplementation, *in vivo* measurement of SCFA production, and *in vitro* tests of microbiota metabolic activity. It would also be useful for such studies to explore the impact of prebiotic supplementation on host physiology, both *in vitro* and *in vivo*. Specifically, the effects of prebiotic supplementation on colonic epithelial barrier integrity, SCFA receptor (GPR41, 43, 109A) expression, and SCFA transporter (MCT1, SMCT1) expression could provide greater insight into the health impacts of prebiotic supplementation, as well as explain why fecal SCFA concentrations may not mirror the metabolic capacity of gut microbiota.

## Methods

### Cohort

Stool was collected from human donors under a protocol approved by the Duke Health Institutional Review Board (Duke Health IRB Pro00074547) for a prospective longitudinal cohort study and biorepository. Participants whose samples were used in this study were treatment-seeking adolescents with obesity who were newly enrolled in a multi-disciplinary weight management program. All subjects received family-based intensive lifestyle modification. Based on clinical necessity, some participants also were placed on a low-carbohydrate diet, medications to facilitate weight loss, or underwent weight loss surgery (Table S2). Due to the low number of patients assigned to each treatment arm, we did not attempt to base any analyses on patient treatment plan. Patients were aged 10-18, with BMI ≥ 95^th^ percentile. None had prior antibiotic use in the 1 month prior to enrolment, used medications known to interfere with the intestinal microbiome, and did not have other significant medical problems. Stool samples used in this study were from enrollment, 3-month, 4.5-month, and 6-month follow-up visits (Table S2). The clinical metadata used for correlations was collected at enrollment, 3 months, and 6 months. The metadata collected nearest to the stool sample collection date was used in our analyses.

### Stool Collection

Patients collected intact stool samples in the clinic or at home using a plastic stool collection container (Fisher Scientific: 02-544-208) and were asked to immediately store this container in their home freezer. Patients then returned the sample by either bringing it to the study team or scheduling a home pickup within 18 hours of stooling. Stool was transported frozen in an insulated container with an ice pack. Upon receipt in the lab, samples were placed on dry ice until transferred to a −80°C freezer for long term storage. All patient samples were frozen at −80°C within 19 hours of stooling (range, 0.08hr – 18.83hr; median, 11.42hr) except one which was stored 44.03hr after stooling. The time between stooling and freezing at −80°C did not have a significant effect on average SCFA production (p = 0.58, ρ = −0.15, Pearson correlation). Stool samples for analysis were processed by removing containers from −80°C storage and thawing on ice in a biological safety cabinet until soft enough to aliquot. Thawed containers of stool were opened to atmosphere for a maximum of 10 minutes while samples were aliquoted. After primary aliquoting, the remaining stool was transferred to an anaerobic chamber (COY Laboratory Products, 5% hydrogen, 5% CO2, 90% Nitrogen) and further portioned into approximately 2g aliquots for this study. These aliquots were then stored as solid stool pellets at −80°C until used for this study.

### *In vitro* fermentation

See figure 1 for an overview of *in vitro* fermentation methods. Aliquoted stool was thawed at room temperature in an anaerobic chamber. Once thawed, stool was weighed and placed into a polyethylene filter bag with 0.33mm pore size (Whirl-Pak B01385) and 10mL of anaerobic 1X PBS was added for each gram of stool, resulting in a 10% w/v fecal slurry, similar to previous studies (41, 42, 70, 71). During our validation experiments, a medium designed to simulate colonic contents was used in place of 1X PBS to create stool slurries (46). The filter bag was then closed and placed into a stomacher (Seward Stomacher 80) where the contents were homogenized on the medium speed setting for 60 seconds. The liquid fraction was removed from the downstream side of the filter membrane, and the solid fraction was discarded. A 1mL aliquot of this liquid fraction was removed for analysis of SCFA concentration, to determine the SCFA concentration of the starting stool sample. During our validation experiments, two separate 1mL aliquots of this liquid fraction were removed: one was used to estimate relative bacteria abundance of starting fecal slurries using total DNA, as has been previously published (72); the remaining aliquot was used to determine the pH of the starting fecal slurry using a handheld pH meter (Elite pH Spear, Thermo-Fischer Scientific). The remaining liquid fraction was incubated in duplicate across six different treatments, either supplemented with inulin (Now Foods Inulin Powder, part #2944), fructooligosaccharides (FOS; Cargill, part #100047199), galactooligosaccharides (GOS; Bimuno Powder), xylooligosaccharides (XOS; BioNutrition prebiotic with Llife-Oligo, part #359), wheat dextrin (Benefiber Original), or unsupplemented. For each reaction, 1mL of 10% fecal slurry was placed in one well of a 24-well cell culture plate. Each well was then delivered 1mL of 1% (w/v) prebiotic solution in 1X PBS; or 1mL of 1X PBS without prebiotic. During our validation experiments, prebiotics were dissolved in colonic medium instead of 1X PBS. The resulting fermentation conditions where therefore 5% fecal slurry with 0.5% prebiotic (w/v). A 5% fecal slurry was selected because its fermentative capacity has been previously demonstrated to be insensitive to small variations in concentration and is feasible to work with using this method (42). A 0.5% final concentration of prebiotic in the context of a 5% fecal slurry is analogous to an average adult consuming 20g of dietary fiber per day, assuming an average daily stool mass of 200g (73). Fermentation reactions were carried out in an anaerobic chamber at 37°C for 24 hours. Following fermentation, 1mL media was taken from each reaction vessel for SCFA quantification. During our validation experiments, a separate 1mL aliquot was taken for pH measurement.

### Simulation of Freeze/Thaws Experienced by Study Samples

To test the effects of freeze/thaw cycles on *in vitro* SCFA production, we collected fresh, whole fecal samples from four healthy adults who were not patients in the study cohort. Informed consent was obtained from volunteers and the protocol was approved by the Duke Health Institutional Review Board. Samples were brought into an anaerobic chamber after voiding. Once in anaerobic conditions, these samples were divided into three aliquots. One aliquot was processed immediately following the same *in vitro* fermentation protocol used in our study. transferred to −80C storage. After a minimum of 24 hours, one of these two aliquots was removed from the freezer and thawed at room temperature for 2 hours, before being returned to −80C for an additional minimum of 24 hours. Each of these frozen aliquots was thawed and processed following the same *in vitro* fermentation protocol. This allowed direct comparison of samples that had been used in fermentations immediately after voiding to those that had been frozen and thawed one and two times.

### Media Preparation

To validate our methods, namely our use of a 5% fecal slurry in PBS, without supplementation of other nutrient components, we compared SCFA production with our methods to SCFA production when stool was instead resuspended in a medium designed to simulate the large intestine. We used a slightly modified medium derived from Gamage et al. 2017 (46). The media contained, per liter: peptone 0.5g, yeast extract 0.5g, NaHCO3 6g, hemin solution (0.5% (w/v) hemin and 0.2% (w/v) NaOH) 100uL, L-cysteine HCl monohydrate 0.53g, bile salts 0.5g, Vitamin Supplement (ATCC MD-VS) 1mL, K2HPO4 0.228g, KH2PO4 0.228g, (NH4)2SO4 0.228g, NaCl 0.456g, MgSO4 0.0456g, CaCl2 0.0460g, Trace Mineral Supplement (ATCC MD-TMS) 1mL, and 287uL glacial acetic acid. The pH of the medium was adjusted to 7.0±0.1.

### Quantification of SCFA

The SCFA concentration of fecal slurries and fermentation vessels was determined following a protocol adapted from Zhao, Nyman, and Jönsson (74). First, a 1mL aliquot of either 10% fecal slurry in PBS or the fermentation vessel contents was obtained. To this, 50 μL of 6N HCl was added to acidify the solution to a pH below 3. The mixture was vortexed, centrifuged at 14,000rcf for 5 minutes at 4°C to remove particles. Avoiding the pellet, 750 μL of this supernatant was passed through a 0.22μm spin column filter. The resulting filtrate was then transferred to a glass autosampler vial (VWR part #66009-882).

Filtrates were analyzed on an Agilent 7890b gas chromatograph (GC) equipped with a flame-ionization detector (FID) and an Agilent HP-FFAP free fatty-acid column (25m x .2mm id x .3μm film). A volume of 0.5μL of the filtrate was injected into a sampling port heated to 220°C and equipped with a split injection liner. The column temperature was maintained at 120°C for 1 minute, then ramped to 170°C at a rate of 10°C/min, then maintained at 170°C for 1 minute. The helium carrier gas was run at a constant flow rate of 1mL/min, giving an average velocity of 35 cm/sec. After each sample, we ran a one minute post-run at 220°C and a carrier gas flow rate of 1mL/min to clear any residual sample. All C2:C5 short-chain fatty acids were identified and quantified in each sample by comparing to an 8-point standard curve that encompassed the sample concentration range. Standards contained 0.1mM, 0.2mM, 0.5mM, 1mM, 2mM, 4mM, 8mM, and 16mM concentrations of each SCFA.

### DNA Extraction, PCR Amplification, and Sequencing

We performed 16S rRNA gene amplicon sequencing on human stool samples to determine microbiota community composition. DNA was extracted from frozen fecal samples with the Qiagen DNeasy PowerSoil DNA extraction kit (ID 12888-100). Amplicon sequencing was performed using custom barcoded primers targeting the V4 region of the 16S gene (75), using published protocols (75–77). The sequencing library was diluted to a 10nM concentration and sequenced using an Illumina MiniSeq and a MiniSeq Mid Output Kit (FC420-1004) with paired-end 150bp reads.

### Identifying Sequence Variants and Taxonomy Assignment

We used an analysis pipeline with DADA2 (78) to identify and quantify sequence variants, as previously published by Silverman et al. (79). To prepare data for denoising with DADA2, 16S rRNA primer sequences were trimmed from paired sequencing reads using Trimmomatic v0.36 without quality filtering (80). Barcodes corresponding to reads that were dropped during trimming were removed using a custom python script. Reads were demultiplexed without quality filtering using python scripts provided with Qiime v1.9 (81). Bases between positions 10 and 150 were retained for the forward reads and between positions 0 and 140 were retained for the reverse reads. This trimming, as well as minimal quality filtering of the demultiplexed reads was performed using the function fastqPairedFilter provided with the DADA2 R package (v1.8.0). Sequence variants were inferred by DADA2 independently for the forward and reverse reads of each of the two sequencing runs using error profiles learned from all 20 samples. Forward and reverse reads were merged. Bimeras were removed using the function removeBimeraDenovo with default settings. Taxonomy was assigned using the function assignTaxonomy from DADA2, trained using version 123 of the Silva database.

### Modeling Microbial Composition Data

To associate microbial genera to SCFA production on different prebiotics, the sequence variant table was amalgamated to the genus level using the R package *phyloseq* (81). Genera that were observed with at least 3 counts in at least 3 samples were retained. This filtering step retained 99.3% of sequence variant counts and a total of 97 genera.

To associate microbial composition to SCFA production on different prebiotics we made use of Bayesian Multinomial Logistic-Normal linear regression implemented in the R package *stray* as the function *pibble* (82). We chose this method to account for uncertainty due to counting, and compositional constraints as motivated in Silverman, Durand (79) and Grantham, Reich (83). Our regression model was defined for the *j*-th sample by the covariate vector *x_j_ =* [1, *x_j_*_(*insulin*)_, *x_j_*_(*GOS*)_, *x_j_*_(*XOS*)_, *x_j_*_(*Dextrin*)_]^*T*^ where *x_j_*_(*insulin*)_ is the amount of total SCFA produced by the community in sample *j* as assessed by our *in vitro* assay and the preceding 1 represents a constant intercept. The regression model priors required that 4 hyperparameters *Gamma*, *Theta, Xi,* and *upsilon* be specified. We set the hyperparameters *Theta* to an *D × Q* matrix of zeros (where *D = 97*, the number of sequence variants; and *Q = 5*, the number of covariates) representing our prior assumption that, on average, the association between each prebiotic and each taxon is zero.

We set the hyperparameters *Gamma* to be the matrix

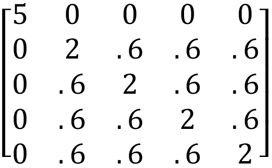

which was chosen to reflect the following prior information: (1) the relative scale of *Gamma_11_* to *Gamma_kk_* (for *k* ∈ ~2, …, 5}) implies that we have little knowledge regarding the mean composition between individuals but that we conservatively expect that the association between butyrate production and microbial composition is small comparatively. (2) the value of 0.6 state that, on average across genera, we assume that the effects each prebiotic are correlated with an average correlation of 0.3, (3) in concert with our prior choices for *Xi* and *upsilon* (below), the scale of *Gamma* represents our assumption that the technical noise in our community measurements is smaller (by a factor of *≈ e*^2^) than the magnitude of the biological variation between samples. This later prior regarding technical versus biological variation was informed by Silverman, Durand (79). All prior choices were further investigated using prior predictive checks (84). To reflect a weak prior assumption that the absolute abundance of each taxon is uncorrelated we choose *upsilon = D + 3* and *Xi* to be the *(D − 1) × (D − 1)* matrix with elements *Xi_ii_ = (upsilon − D)* and *Xi_ij_ = (upsilon − D)/2*, for *j ≠ i* (85). While the model fit by *stray* and our corresponding priors were specified with respect to additive log-ratio coordinates, we utilized theory from compositional data analysis to transform these results into centered log-ratio coordinates for interpretation (79). Credible intervals and figures reflect 2000 samples from the posterior distribution of the corresponding multivariate regression model.

## Acknowledgements

This work was supported in part by NIH grants R24-DK110492 and 1R01DK116187, the Translational Research Institute through Cooperative Agreement NNX16AO69A, the Damon Runyon Cancer Research Foundation, and the UNC CGIBD (NIDDK P30DK034987). This work used a high-performance computing facility partially supported by grant 2016-IDG-1013 (“HARDAC+: Reproducible HPC for Next-generation Genomics”) from the North Carolina Biotechnology Center. The authors would also like to acknowledge Jessica McCann, Cameron Catherine, Charles Sarria, Alexandra Zizzi, and Janet Wooton for their contributions to this work.

## SUPPLEMENTARY FIGURES AND TABLES

**Figure S1:**
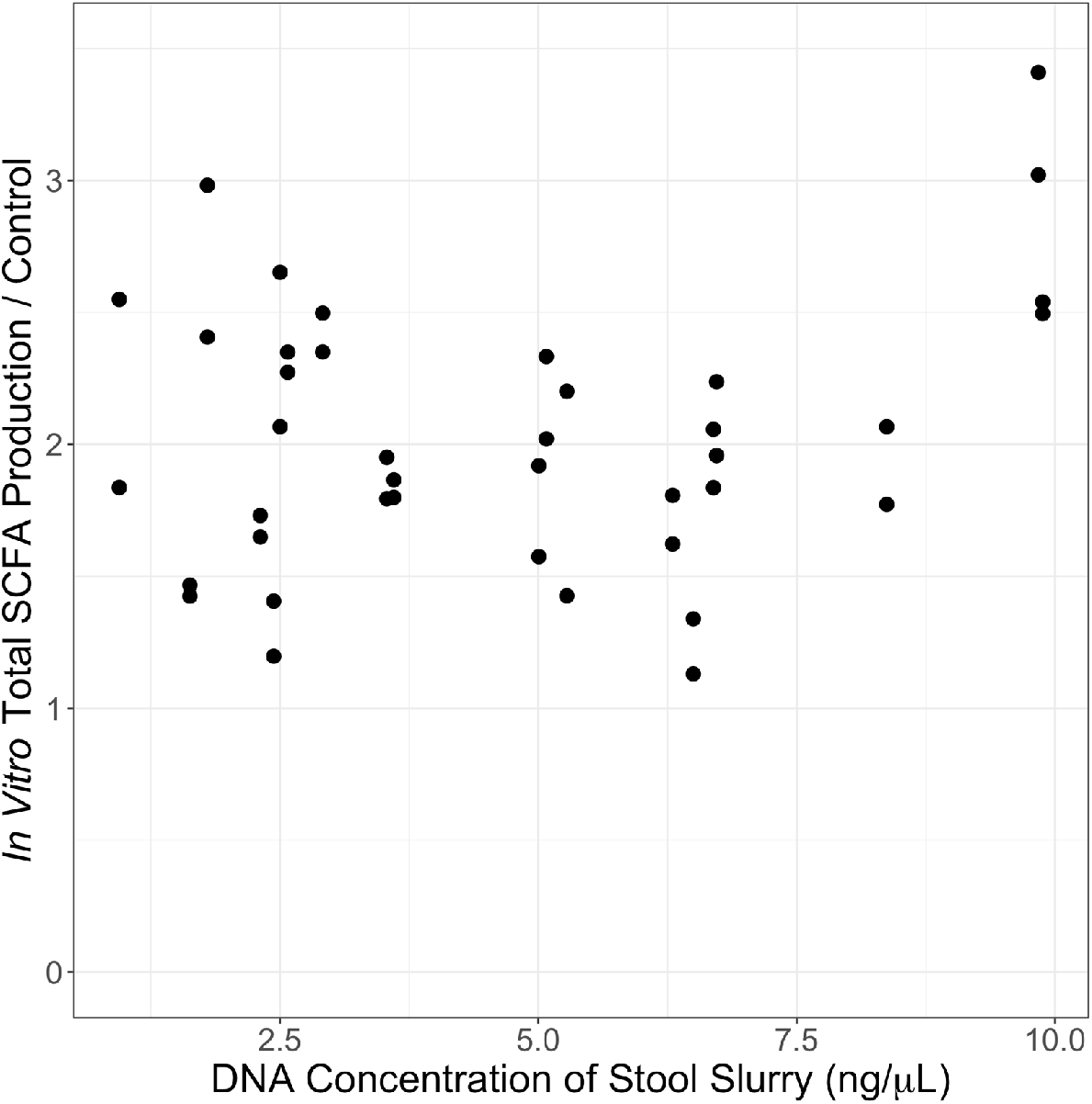
Bacteria abundance in stool, as measured by DNA concentration, does not correlate with control-corrected total SCFA production in *in vitro* cultures (p = 0.38, ρ = 0.14; Spearman correlation).

**Figure S2:**
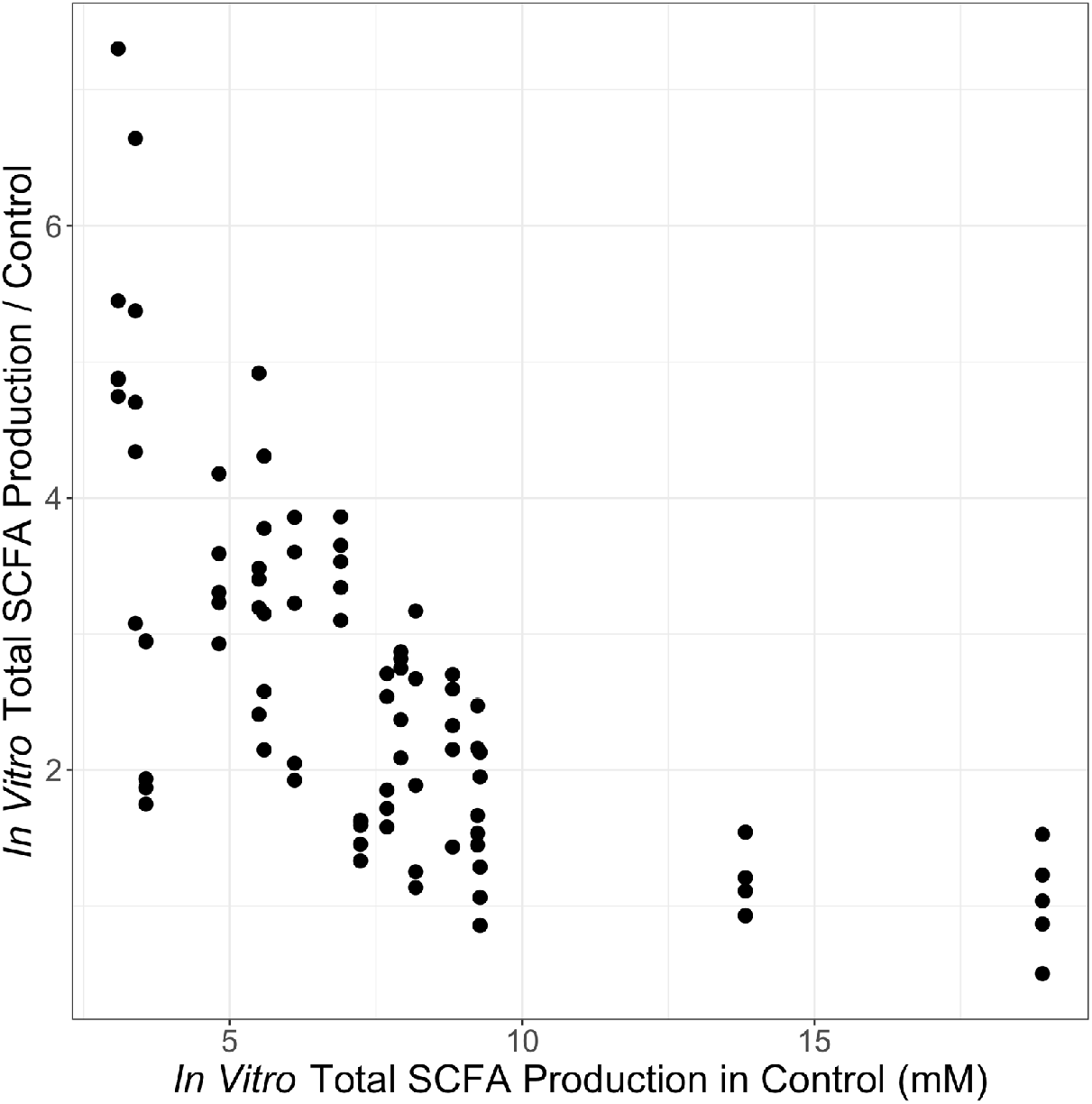
To control for differences in overall cell viability or stool slurry nutrient content between donors, we corrected measurements of SCFA concentration by dividing the treatment SCFA concentration by the control SCFA concentration. The resulting fold-change data do not contain information about absolute SCFA production. We examined the potential for this artifact to influence our interpretation, and found that fold changes of SCFA concentrations after prebiotic treatment relative to the control were correlated with absolute control treatment levels (p<0.0001, ρ = −0.77, Spearman correlation), Absolute (not corrected to control) SCFA concentrations are presented in figure S3.

**Figure S3:**
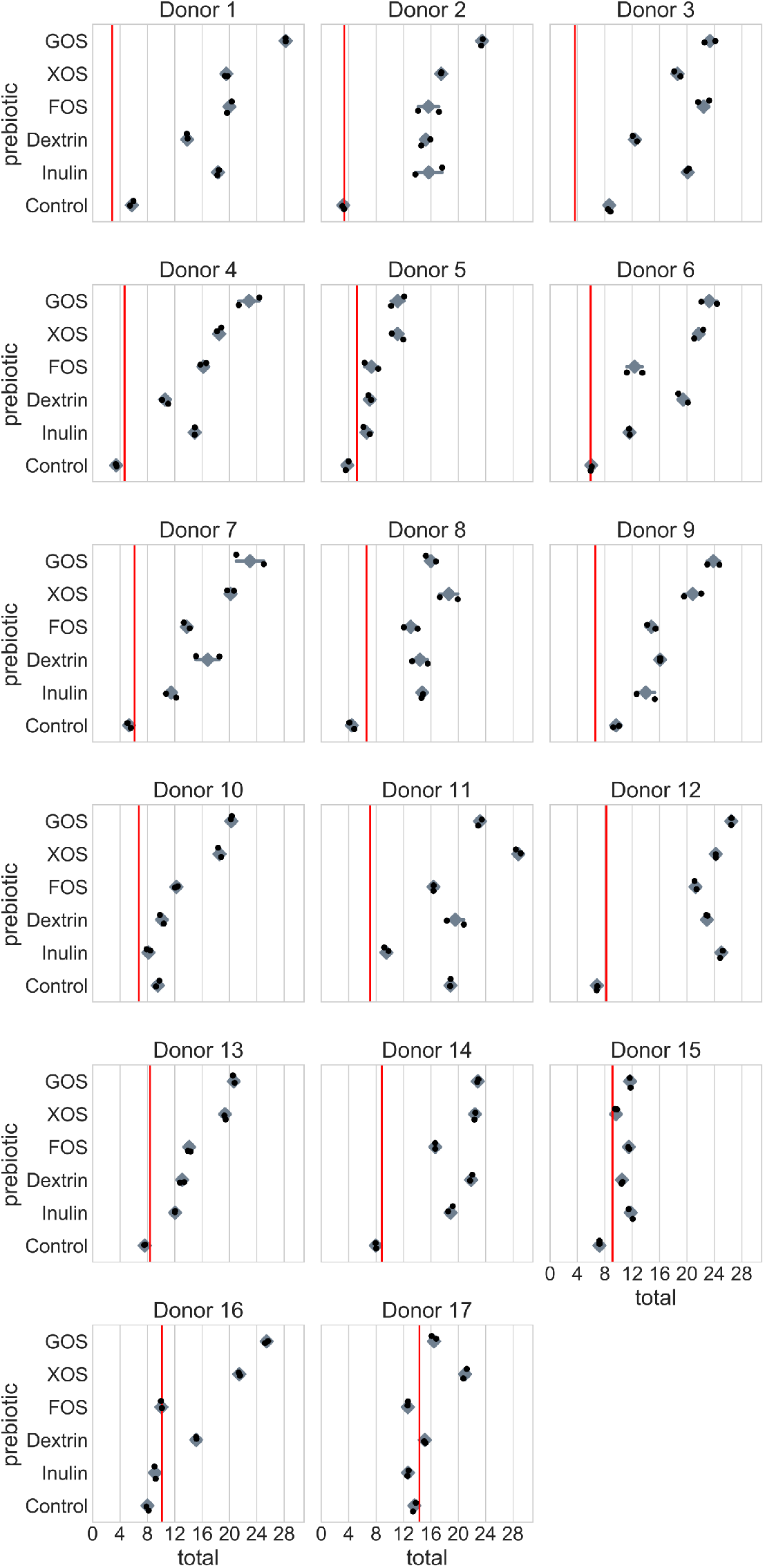
Total SCFA concentration of *in vitro* fermentation vessels after 24hr fermentation, plotted for each donor across five prebiotic treatments and the unsupplemented control vessel. The vertical red line indicates the total SCFA concentration of the starting fecal slurry prior to fermentation. Instances where SCFA concentration decreases during fermentation may be explained by net SCFA consumption by the community when no fermentable carbon is supplied, or by a lack of change in concentration coupled with technical variation in our measurements. Instances where SCFA concentration is increased in the control treatment suggest that some unmetabolized carbohydrate may have remained in the stool to be metabolized during *in vitro* fermentation.

**Figure S4:**
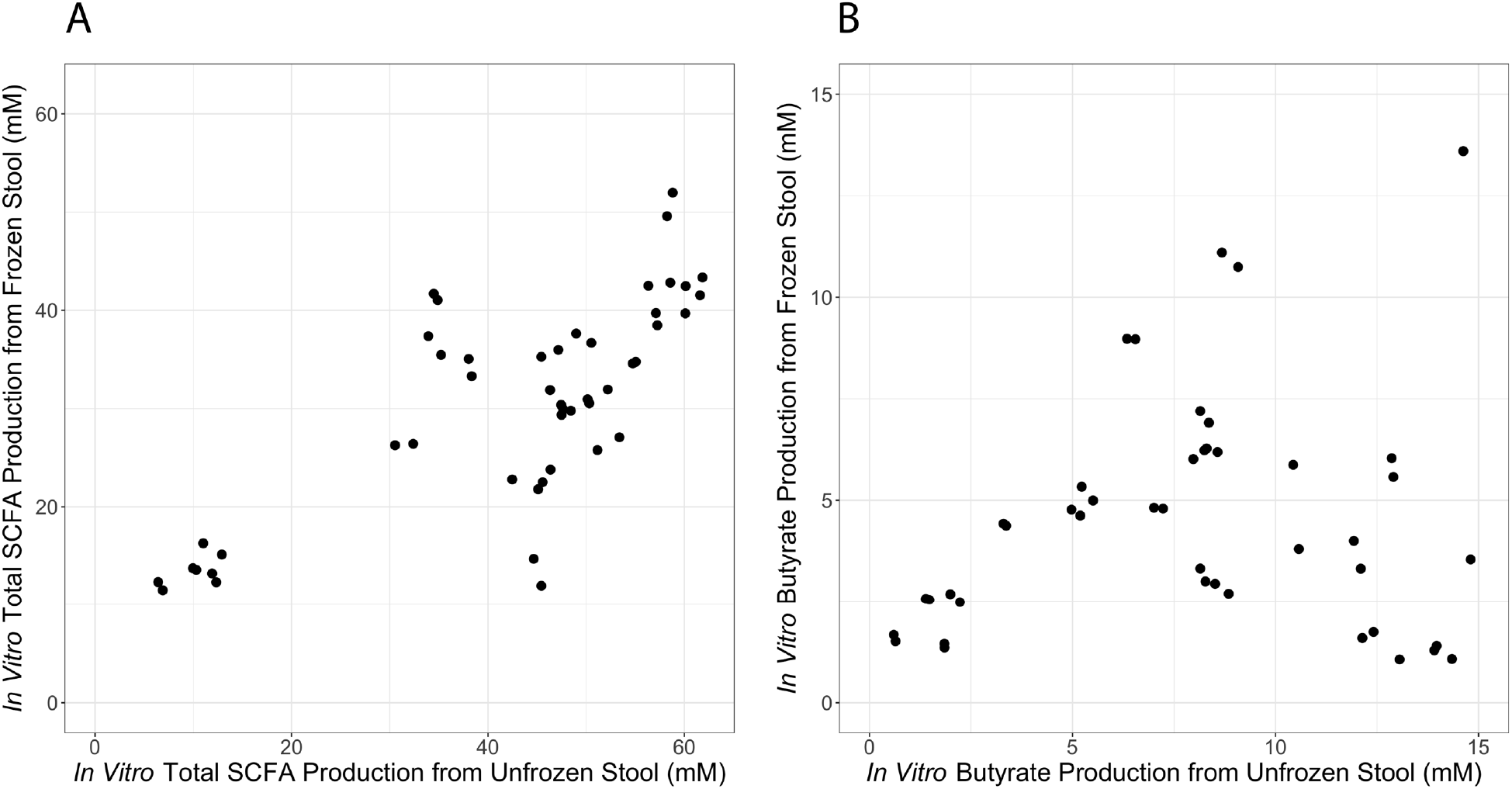
*In vitro* total SCFA production from unfrozen stool samples and from twice frozen stool samples is highly correlated (A, p < 0.0001, ρ = 0.75, Spearman correlation). In contrast, *in vitro* butyrate production is not correlated between unfrozen and twice frozen stool samples (B, p = 0.18, ρ = 0.19 Spearman correlation).

**Figure S5:**
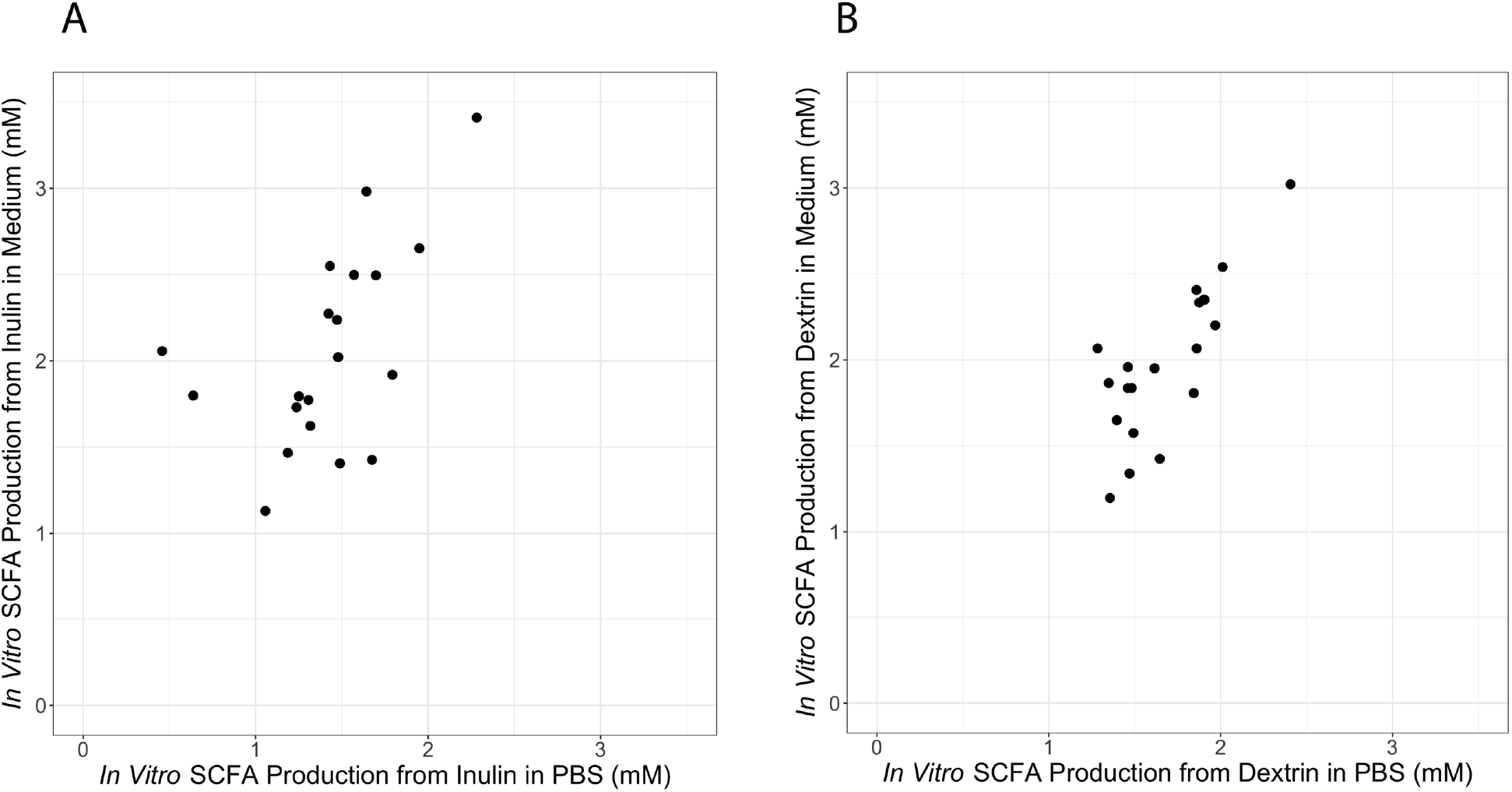
*In vitro* total SCFA production from inulin (A) and dextrin (B) is well correlated between cultures grown in PBS and cultures grown in a medium designed to mimic the colonic environment (p = 0.001, ρ = 0.68 (A); p = 0.02, ρ = 0.51 (B); Spearman correlations).

**Figure S6:**
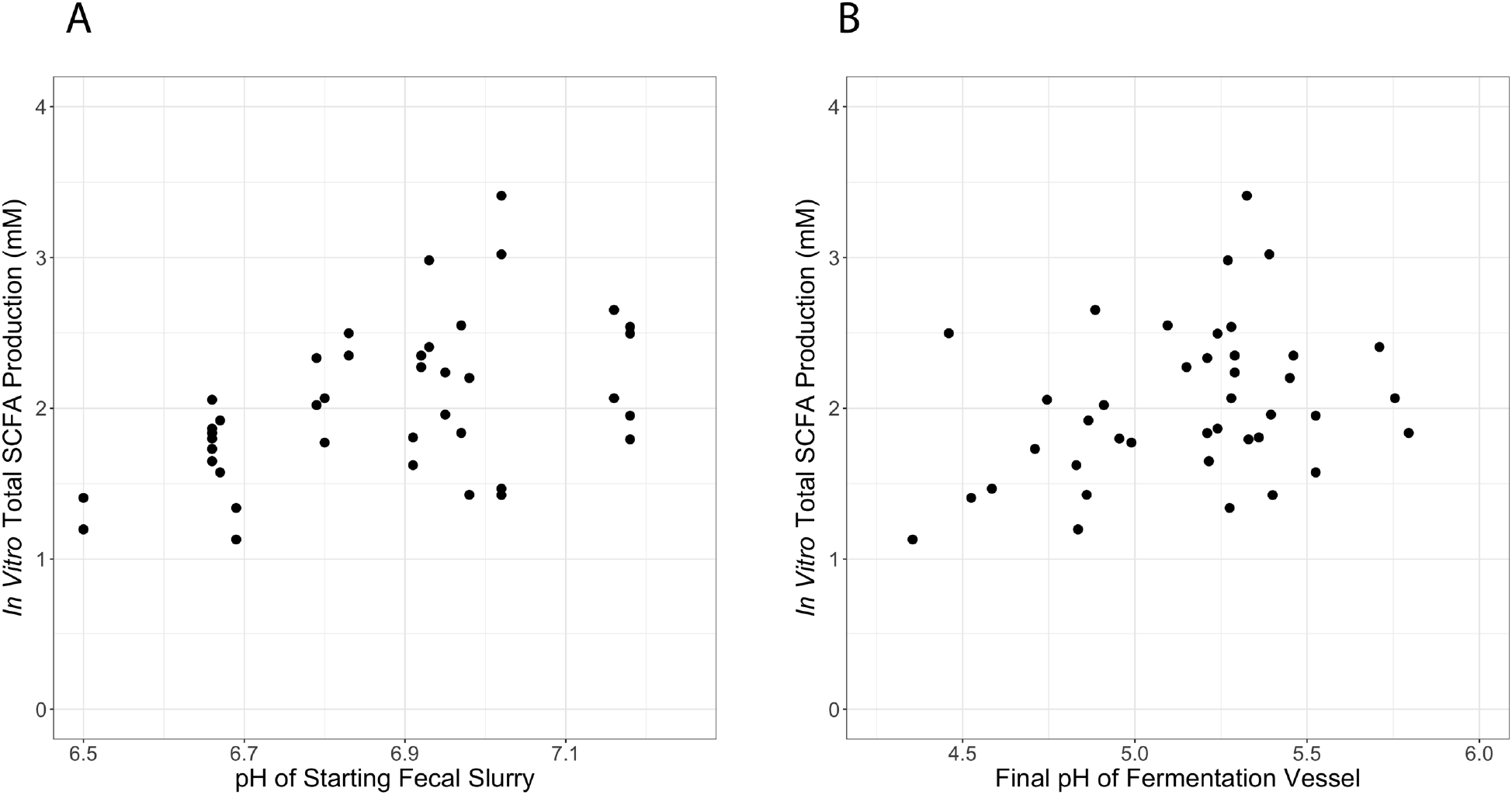
*In vitro* total SCFA production over control is positively correlated with the pH of starting fecal slurries (p = 0.003, ρ = 0.46; Spearman correlation). A weaker correlation might exist between SCFA production and the final pH of fermentation vessels (p = 0.067, ρ = 0.29; Spearman correlation).

**Figure S7:**
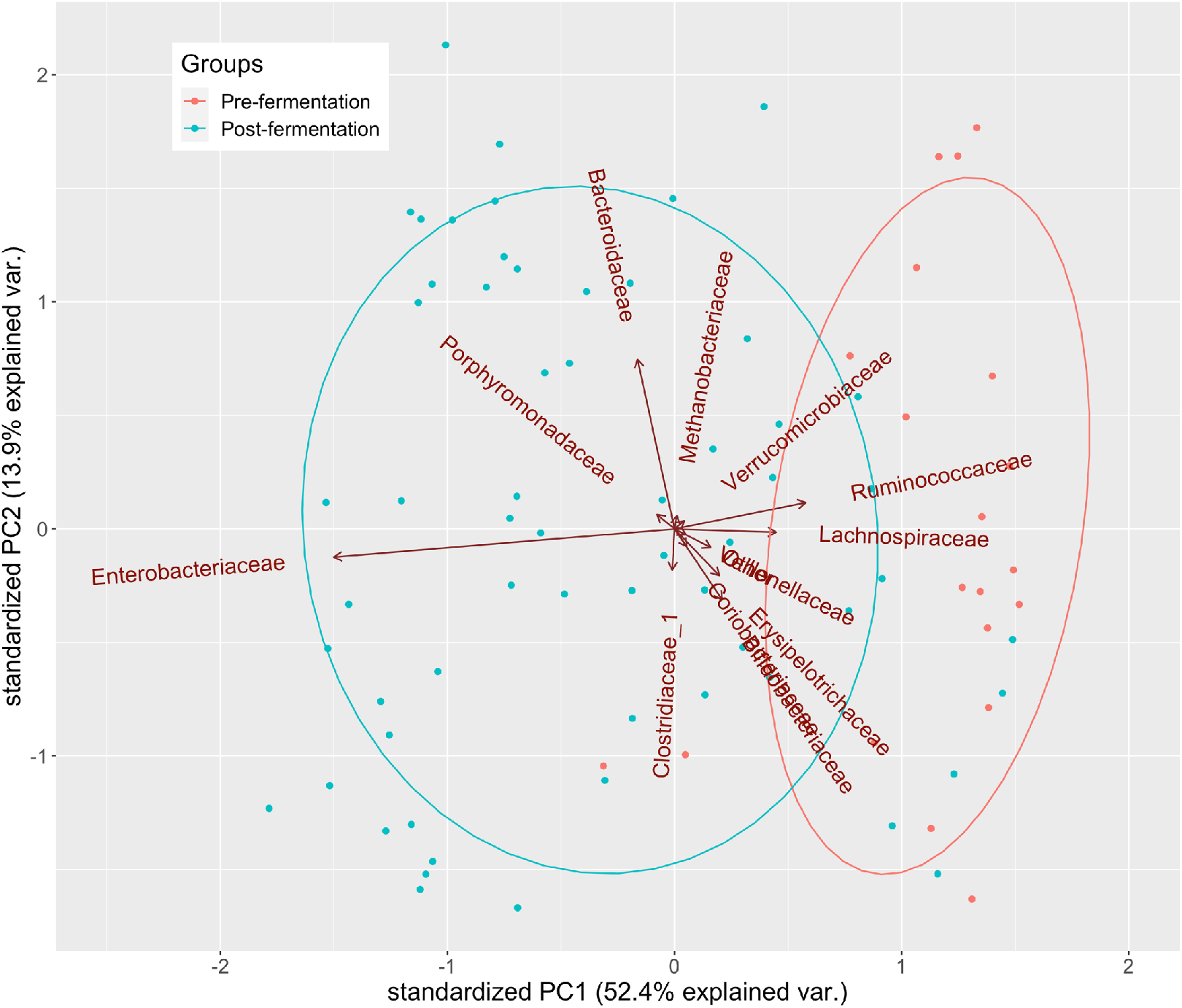
Microbial community composition changes during the course of a 24hr fermentation. We performed 16S rRNA sequencing on pre-fermentation and post-fermentation samples from 20 donors across 3 treatments (inulin, dextrin, and control). We then measured the Shannon diversity of the fecal slurries both before and after fermentation (all treatments averaged) and found a significant decrease in Shannon diversity over the course of fermentation (p < 0.0001; paired t-test). To characterize the changes in community composition associated with this decrease in diversity, we tested the pre-fermentation and post-fermentation samples for differential abundance of taxa at the species level. We found 10 taxa with significantly different abundances between the two sample sets (p < 0.05; Benjamini-Hochberg corrected Wilcoxon rank-sum tests). Of these 10 taxa, only Escherichia/Shigella spp. increased in relative abundance after fermentation, while nine other taxa each decreased in relative abundance (*Anaerostipes hadrus, Bacteroides acidifaciens, Blautia faecis, Blautia wexlerae* and two other *Blautia* of undetermined species, two undetermined species in the *Eubacterium hallii* group, and *Ruminococcaceae_UCG-004 spp*.). We attribute these changes to differences in growth rates among bacteria in our in vitro system, and the inability of our fermentation medium to support the growth of some community members.

**Table S1:**
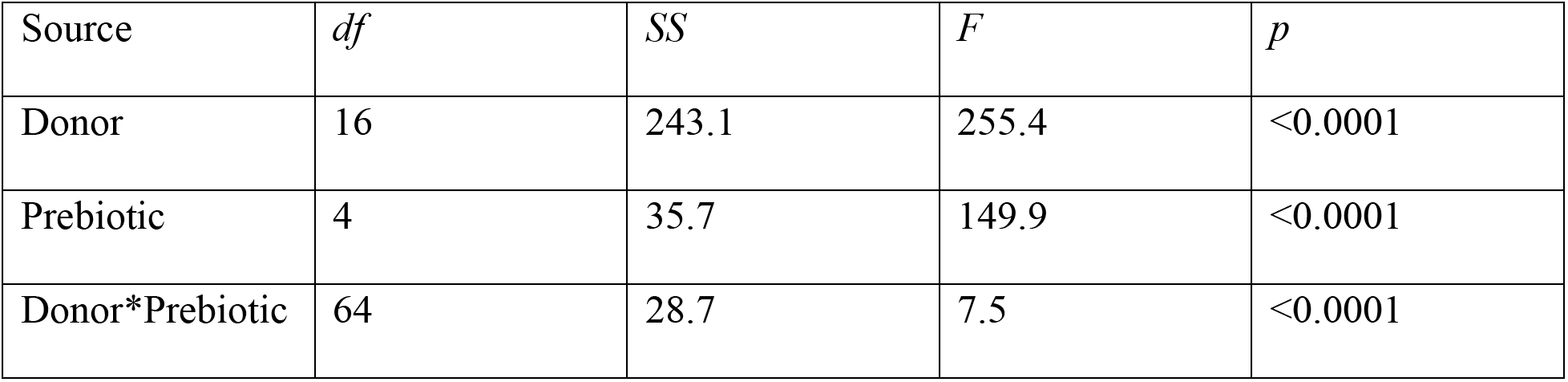
Two-way ANOVA of *in vitro* SCFA production across donors and prebiotics.

| Source | *df* | *SS* | *F* | *p* |
| --- | --- | --- | --- | --- |
| Donor | 16 | 243.1 | 255.4 | <0.0001 |
| Prebiotic | 4 | 35.7 | 149.9 | <0.0001 |
| Donor*Prebiotic | 64 | 28.7 | 7.5 | <0.0001 |

**Table S2:** Treatment arms and sample timepoints of patients in this study.

| Patient | Treatment | Timepoint (month) |
| --- | --- | --- |
| 1 | Lifestyle only | 6 |
| 2 | Lifestyle only | 0 |
| 3 | Lifestyle only | 3 |
| 4 | Lifestyle only | 0 |
| 5 | Lifestyle only | 6 |
| 6 | Lifestyle only | 4.5 |
| 7 | Lifestyle only | 0 |
| 8 | Lifestyle only | 3 |
| 9 | Lost to follow-up | 0 |
| 10 | Lifestyle only | 4.5 |
| 11 | Lifestyle only | 3 |
| 12 | Lifestyle only | 4.5 |
| 13 | Low carb | 0 |
| 14 | Lifestyle only | 4.5 |
| 15 | Metformin | 4.5 |
| 16 | Lifestyle only | 4.5 |
| 17 | Lifestyle only | 4.5 |

